# Spatiotemporal contribution of neuromesodermal progenitor-derived neural cells in the elongation of developing mouse spinal cord

**DOI:** 10.1101/2020.05.03.075382

**Authors:** Mohammed R Shaker, Ju-Hyun Lee, Kyung Hyun Kim, Veronica Jihyun Kim, Joo Yeon Kim, Ji Yeoun Lee, Woong Sun

## Abstract

During vertebrate development, the posterior end of the embryo progressively elongates in a head-to-tail direction to form the body plan. Recent lineage tracing experiments revealed that bi-potent progenitors, called neuromesodermal progenitors (NMPs), produce caudal neural and mesodermal tissues during axial elongation. However, their precise location and contribution to spinal cord development remain elusive. Here we used NMP-specific markers (Sox2 and BraT) and a genetic lineage tracing system to localize NMP progeny *in vivo*. NMPs were initially located at the tail tip, but were later found in the caudal neural tube, which is a unique feature of mouse development. In the neural tube, they produced neural stem cells (NSCs) and contributed to the spinal cord gradually along the AP axis during axial elongation. Interestingly, NMP-derived NSCs preferentially contributed to the ventral side first and later to the dorsal side at the lumbar spinal cord level, which may be associated with atypical junctional neurulation in mice. Our current observations detail the contribution of NMP progeny to spinal cord elongation and provide insights into how different species uniquely execute caudal morphogenesis.

## INTRODUCTION

In vertebrates, neuroepithelial cells give rise to the neural tube, which forms through two processes along the anterior-posterior (AP) axis. The first process is primary neurulation, which progresses through convergent extension, elevation, bending, and fusion of the neural plate, forming the rostral neural tube (Pai et al., 2012). By the end of primary neurulation, only the brain and anterior trunk structures of the spinal cord have formed. As the embryo develops, progressive addition of new neural stem cells (NSCs) is required at the posterior end of the spinal neural tube for neural tube elongation. The cells at the dorsal region of the tail bud aggregate and the tail bud ultimately undergoes cavitation, forming the caudal neural tube; this comprises the future caudal domain of the spinal cord, which is in continuity with the neural tube in the trunk derived from the primary neurulation (Schoenwolf, 1984). Defects in this process, termed “secondary neurulation,” are often associated with spina bifida, a common congenital malformation in humans (Yang et al., 2014). Secondary neurulation is an embryonic process contributing to axial neural elongation; thus, here we use the term “neural tube elongation” instead of “secondary neurulation.” Vertebrates exhibit morphological differences during neural tube elongation. For instance, chick and human tail bud cells undergo cavitation to generate the elongating neural tube; eventually, the elongating neural tube adheres to the primary neural tube at the junctional neurulation zone that is clearly present in the chick and humans (Dady et al., 2014; Saitsu et al., 2004), but not in mouse (Schoenwolf, 1984). In the mouse, elongating neural tube cells are polarized via cell rearrangement, and later migrate toward the lumen of the primary neural tube and fuse, establishing a continuous neural tube without a junctional zone (Colas and Schoenwolf, 2001). In addition, the chick neural tube elongation process shapes the spinal cord and extends to the lumbar region (Colas and Schoenwolf, 2001). A similar anatomical feature is also observed in humans (O’Rahilly and Müller, 2003; Saitsu et al., 2004), which is contrary to that observed in mice, in which only the tail is formed by neural tube elongation (Shum et al., 2010).

Another important aspect of caudal tube elongation is that the NSCs at this level are derived from neuromesodermal progenitor cells (NMPs), which are able to produce both neural and mesodermal tissues (Tsakiridis and Wilson, 2015; Tzouanacou et al., 2009). NMPs are bipotent progenitor cells that co-express Sox2 and Brachyury T (BraT), which exist in the tail- bud of human, chick (Olivera-Martinez et al., 2012), and mouse embryos (Anderson et al., 2013). NMPs persist over extended periods and are eventually depleted toward the end of body axis elongation (Wymeersch et al., 2016). In addition, fate mapping studies at the tail bud stage using TCreER2 transgenic mice further identified BraT-positive cells located in the floor plate of the neural tube (Anderson et al., 2013). Collectively, this indicates that anterior and posterior neural tissues are formed from independent lineages of cells, which challenges the traditional concept of segregation of the three germ layers (ectoderm, endoderm, and mesoderm) and subsequent neural fate assignment in ectodermal tissues. While it is well described that NMPs contribute to spinal cord elongation, quantitative and comprehensive analyses into what extent NMPs contribute to spinal cord development are currently missing.

In this study, we present detailed descriptions of the spatiotemporal localization of NMPs during the axial elongation periods in the mouse and chick embryos and show that NMPs localize within mouse neural folding and elongation tissues, whereas chick NMPs only localize at the tail bud tissue. Furthermore, we mapped the distribution and neuronal differentiation of NMP-derived NSCs depending on their birthdates using a genetic BraT lineage reporter system. Our current observations clearly illustrate that NSCs derived from NMPs contribute to the caudal spinal cord with developmental orders depending on the anteroposterior and dorsoventral (DV) axes.

## RESULTS

### Presence of neuromesodermal progenitors in the mouse neural tube

NMPs emerge near the primitive streak and later remain in restricted regions in the tail bud during axial elongation (Wymeersch et al., 2016). We extended the observation of the spatiotemporal distribution of NMPs during different stages of axial elongation of neural tissues, including the early and late stages of neural folding and neural tube elongation. During an early stage of neural folding (E8), whole-mount immunofluorescence labeling revealed that NMPs, which are double-labeled with Sox2 and BraT, were abundant within caudal neuroepithelial sheets before neural folding (Fig. 1A). Furthermore, a population of NMPs was observed within the neural tissues already forming the neural tube (Fig. 1A), suggesting that at least one population of Sox2^+^ cells undergoing neurulation maintains BraT expression. At a later stage of neural folding (E10), these double-positive cells were predominantly found within the neural tube, while tail bud cells prominently expressed only BraT (Fig. 1B). At E11, double-positive cells were detected in a broad domain of the elongating caudal neural tube, while the expression levels of BraT appeared to be diminished in the tail bud (Fig. 1C), and, by E12, double-positive cells were no more observed within the neural tube (Fig. 1D). Cross-sectioning further demonstrated that the neural tube prominently expresses Sox2 only, and that only a few Sox2- and BraT-expressing cells were restricted to the dorsal caudal tail bud domain (Fig. 1D). Considering that the embryonic stage E10 marks the end of the neural folding process and the beginning of neural tube elongation, our data suggest that NMPs are transiently located at the caudal neural tube, even after depletion of NMPs in the tail bud.

**Figure 1.**
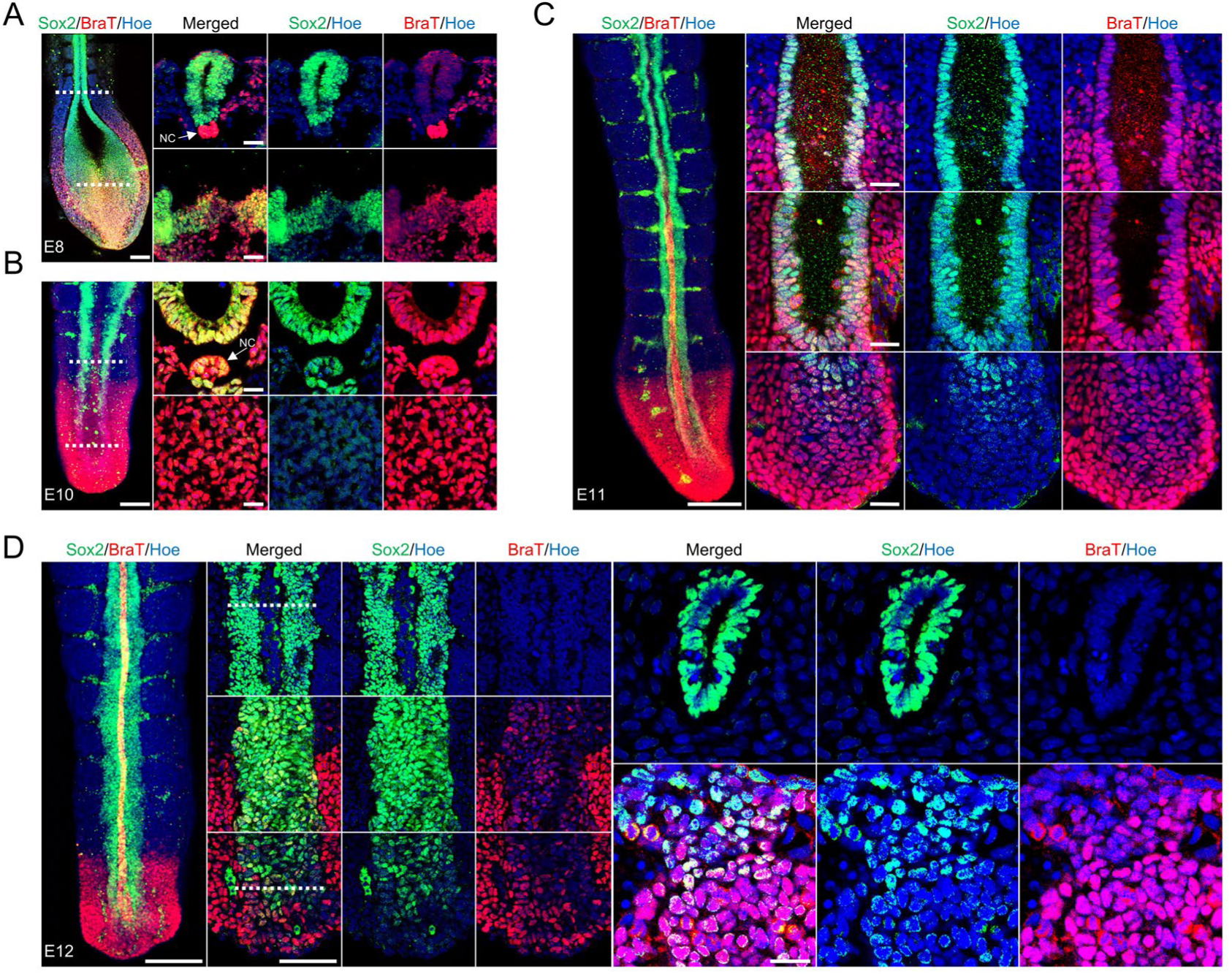
Distribution of neuromesodermal progenitors during mouse neural tube formation and elongation. (A) Dorsal view of whole-mount imaging of E8 caudal neural tube immunostained for Sox2 (Green) and BraT (Red); dotted lines indicate the level of sectioned images. The right panel depicts a transverse section of E8 mouse spinal neural tube during the convergent extension and neural tube formation phases; sections were stained for Sox2 (Green) and BraT (Red). All sections were counterstained with Hoechst 33342 (Blue). Scale bar = 100 µm, scale bar of sectioned images = 40 µm. NC is notochord, white arrow shows the notochord. (B) Dorsal view of E10 whole-mount imaging of a caudal neural tube immunostained for Sox2 (Green) and BraT (Red); dotted lines indicate the level of sectioned images. The right panel depicts a transverse section of E10 caudal spinal cord at tail-bud and neural tube levels, stained for Sox2 (Green) and BraT (Red). All sections were counterstained with Hoechst 33342 (Blue). Scale bar = 100 µm, scale bar of sectioned images = 20 µm. NC is notochord, white arrow shows the notochord. (C) Dorsal view of an E11 tail whole-mount immunostained for Sox2 (Green) and BraT (Red); the right images depict magnified caudal domains of the tail-bud, neural rosette, and elongating neural tube. All sections were counterstained with Hoechst 33342 (Blue). Scale bar = 100 µm, scale bar of magnified images = 20 µm. (D) Dorsal view of whole-mount imaging of an E12 tail immunostained for Sox2 (Green) and BraT (Red); the right images depict magnified caudal domains of tail-bud, neural rosette, and elongating neural tube; dotted lines indicate the level of sectioned images. The right panel depicts the images of a transverse section at the tail-bud and caudal neural tube stained for Sox2 (Green) and BraT (Red). All sections were counterstained with Hoechst 33342 (Blue). Scale bar = 100 µm, scale bar of magnified images = 50 µm, scale bar of sectioned images = 40 µm.

### Absence of neuromesodermal progenitors in the chick neural tube

We described above that double-positive cells, which are potentially NMPs, localize within the mouse spinal caudal neural tube and contribute to the formation and elongation of the caudal neural tube. To determine whether a similar NMP distribution is conserved in the chick, we focused on the events of primary neurulation and neural tube elongation using whole-mount and cross-section analysis. At Hamburger Hamilton (HH) 9, NMPs were found in the tail bud and in mesodermal tissues, with absent BraT expression in neural tissues (Fig. 2A). By HH12, before neural tube elongation starts, NMPs were abundant in the tail bud and ventral clusters beneath the newly forming neural tube (Fig. 2B). However, newly formed primary neural tube expressed Sox2 only, while adjacent notochord and mesodermal tissues maintained BraT expression (Fig. 2B). During chick neural tube elongation, neural cells aggregated at the dorsal tail bud before arranging into a cord-like mass that was continuous with the primary neural tube; these events occurred between HH16 and HH45 (Yang et al., 2003). By HH28, we observed Sox2^+^-cell aggregation at the dorsal tail bud (Fig. 2C). Interestingly, unlike in the mouse, we did not detect NMPs in the chick elongated neural tube; BraT expression was restricted to the midline and the ventral domain of the tail bud and notochord of the newly formed, elongated neural tube and lumbar spinal cord (Fig. 2C), suggesting that NMPs were depleted from the tail bud during axial elongation.

**Figure 2.**
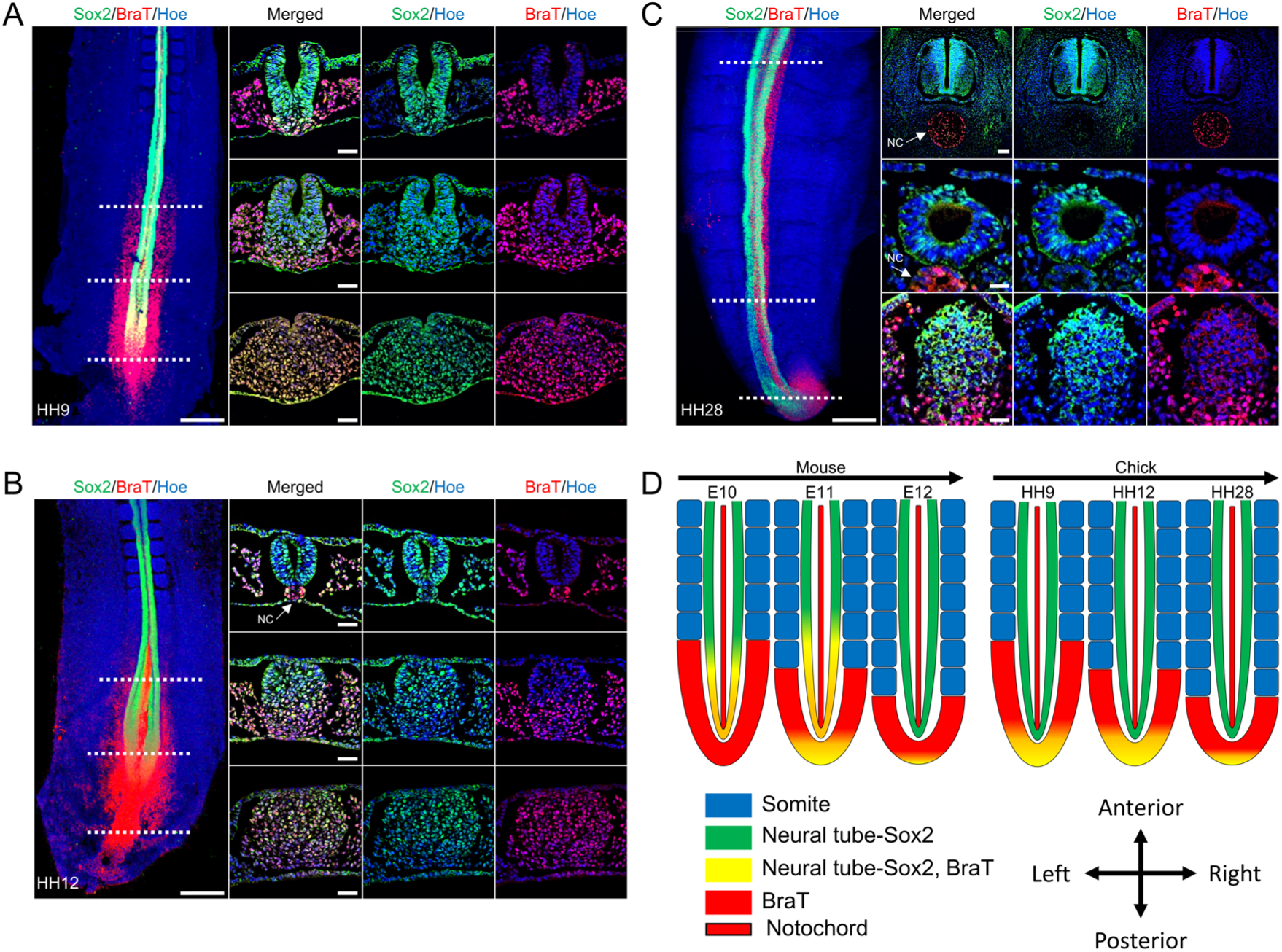
Spatiotemporal expression of neuromesodermal progenitors during chick neural tube formation and elongation. (A) Dorsal view of whole-mount imaging of an HH9 caudal neural tube; dotted lines indicate the level of transverse sectioned images at the tail-bud, neural folding, and elevation and bending levels. All sections were immunostained for Sox2 (Green) and BraT (Red) and counterstained with Hoechst 33342 (Blue). Scale bar = 100 µm, scale bar of sectioned images = 30 µm. (B) Dorsal view of whole-mount imaging of an HH12 caudal neural tube; dotted lines indicate the level of sectioned images at tail-bud, neural rosette, and neural tube level. All sections were immunostained for Sox2 (Green) and BraT (Red) and counterstained with Hoechst 33342 (Blue). Scale bar = 100 µm, scale bar of sectioned images = 30 µm. NC is notochord, white arrow shows the notochord. (C) Dorsal view of whole-mount imaging of an HH28 caudal spinal cord; dotted lines indicate the transverse images of the right panel. All sections were immunostained for Sox2 (Green) and BraT (Red) and counterstained with Hoechst 33342 (blue). Scale bar = 100 µm, scale bar of sectioned images = 30 µm, scale bar of magnified sectioned images = 70 µm. NC is notochord, white arrow shows the notochord. (D) Schematic diagram summarizes the spatiotemporal distribution of NMPs during the phases of neurulation and axial elongation in both the mouse and chick.

In sum, the mouse and chick display distinct morphogenetic processes during neural tube elongation. Our data add additional distinct features where mouse NMPs are included in the elongating neural tube, whereas chick NMPs are restricted to the tail bud and *Sox2*^+^ cells are likely generated only for primary neurulation (Fig. 2D).

### Lineage tracing of NMP-derived NSCs in spinal cord elongation

Multiple clonal studies and single-cell labeling experiments in the mouse and chick have demonstrated the contribution of NMP-derived NSCs to the developing neural tube (Brown and Storey, 2000; Forlani et al., 2003). However, the contribution of NMP-derived cells to later spinal cord development is less addressed. Thus, we explored the fate of NMP-derived NSCs by utilizing transgenic TCreERT2 mice crossed with Rosa-EGFP mice to trace the cells generated from BraT-expressing cells in which Cre recombinase activation is triggered by Tamoxifen (TAM) injection (Anderson et al., 2013). We first traced NMP progeny before neural tube closure by injecting TAM at E5, a stage where blastocysts express BraT at the caudal domain, and gastrulation is being triggered (van den Brink et al., 2014). GFP^+^ cells were expressed throughout the trunk tissues; this expression was maintained during development (Figure 3A), and a detectable number of GFP^+^ cells were observed up to the midbrain and hindbrain domains (Fig. 3A, magnified image). Accordingly, GFP^+^/Sox2^+^ cells were widespread in the entire spinal cord (Fig. 3B).

**Figure 3.**
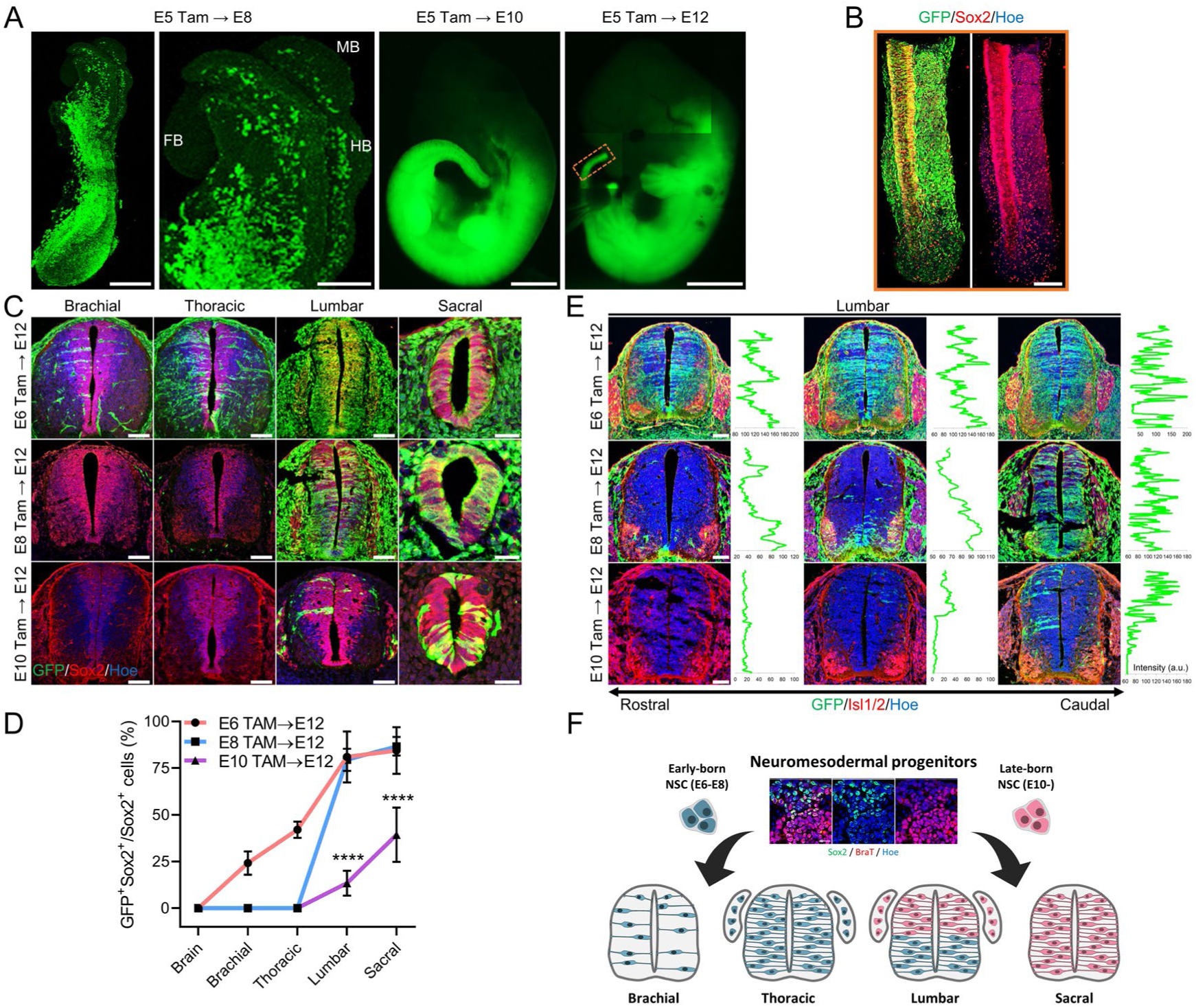
Contribution of neuromesodemral progenitors to the formation of the embryonic central nervous system. (A) Lateral view of E8 mouse embryos treated with tamoxifen at E5 and immunostained for GFP (Green); stained tissue is 3D-imaged using a Leica microscope. E10 and E12 embryos treated with tamoxifen at E5 were imaged by Evos fluorescent microscopy, showing the endogenous expression of GFP immediately after dissection. The orange rectangular box indicates the level of staining in (B). Scale bar = 100 µm, scale bar of E8 magnified anterior domain = 160 µm, scale bar of whole embryos = 2 mm. (B) Lateral-dorsal view of whole-mount E12 tail tissue treated with tamoxifen at E5 and immunostained for Sox2 (Green) and BraT (Red) and counterstained with Hoechst 33342 (Blue). Scale bar = 100 µm. (C) Detection of GFP^+^ cells in the pool of neural progenitors of sectioned embryonic spinal cords derived from three groups of E12 embryos., where first, second, and third groups of embryos treated with tamoxifen at E6, E8, and E10, respectively. Transversely sectioned brachial, thoracic, lumbar, and sacral tissues were immunostained for Sox2 (Green) and BraT (Red) and counterstained with Hoechst 33342 (Blue). Scale bar = 70 µm, scale bar of sacral images = 30 µm. (D) Quantification of GFP and Sox2 double positive cells among Sox2^+^ cells in E6 to E12, E8 to E12, and E10 to E12 tamoxifen-treated embryos depicted in (C). Data are presented as mean ± standard deviation. ^****^ P<0.0001 One Way ANOVA. (E) Transverse section along the rostrocaudal axis of the lumbar region only, immunostained for Sox2 (Green) and Isl1/2 (Red) and counterstained with Hoechst 33342 (Blue). Intensity images depict the dorsal-ventral distribution of GFP within the ventricle of the lumbar neural tube. Scale bar = 70 µm. (F) Schematic diagram illustrating the distribution of GFP^+^ cells derived from NMPs at E6, E8, and E10 along the rostrocaudal level of the lumbar domain. E6 and E8 GFP^+^ cells contribute to the brachial, thoracic, and ventral domain at lumbar level; E10 GFP^+^ cells preferentially generate dorsal, lumbar, and sacral domains.

For detailed analyses, BraT cells were labeled by TAM injection at three different time points, at E6 (before neurulation), E8 (during neurulation), and E10 (during neural tube elongation), and the embryos were harvested at E12 (Fig. 3C, D) to assess the contribution of NMP-derived NSCs ‘born’ at different time points in the embryonic spinal cord. Labeling at E6 led to the emergence of GFP^+^ NSCs from the brachial level (25%), reaching approximately 40% GFP^+^ NSCs within the thoracic neural tube and a progressively greater proportion of NSCs at the caudal parts of the spinal cord (Fig. 3D). On the other hand, tracing of BraT- expressing cell progenitors at E8 illustrated the blunting of brachial and thoracic cell populations and first appeared at the lumbar level. Furthermore, cells derived from E10 NMPs contributed significantly less to lumbar level NSCs, and were more confined to the sacral regions. Collectively, these data indicate that NMP-derived NSCs appear to be sequentially added to the caudal part of the spinal cord during axial elongation.

During our observations, we noticed that there was also a significant tendency for DV preference of NMP-derived NSCs depending on their birthdate. This tendency was most prominent at the lumbar level (Fig. 3E). Labeled NMPs at E8 were preferentially localized to the ventral domain of the rostral lumbar level, while DV-dependent differences were not observed at the lumbosacral level (Fig. 3E). Conversely, labeled NCSs derived from NMPs at E10 were preferentially detected at the dorsal domain of the caudal lumbar neural tube (Fig. 3E). Altogether, these data indicate that NSCs derived from the NMPs at the caudal spinal cord are distributed with AP and DV ordering, as summarized in Fig. 3F.

### The progeny of NMPs generates mature spinal motoneurons and somatic sensory neurons

Because we found that NMP-derived NSCs are located at the ventral and dorsal domains of the spinal neural tube (Fig. 3), we then explored the contributions of these NSCs to the differentiation of corresponding spinal neurons. First, we quantified the percentage of Olig2-labeled motoneuron (MN) progenitors descended from NMPs at the lumbar and sacral levels. MN progenitors derived from NMPs were prominent in lumbar regions, amounting to approximately 60% of MN progenitors born at E6 and E8 (Fig. 4A, B). Similar data were also obtained at the sacral level, with approximately 80% of MN progenitors having derived from NMPs. On the other hand, TAM injection at E10 resulted in an approximate labeling of 20% of cells. Based on the assumption that lineage labeling gave rise to the above cumulative picture owing to the labeling of later-stage NMPs as well as NSCs derived at the timepoint of TAM injection, we estimated the proportion of MN progenitors that descended from NMPs at different durations of labeling (Fig. 4C). By this estimation, it appeared that 60% of lumbar MN progenitors and 74.7% of sacral MN progenitors were born during the E8-E10 period, while 8.5% of lumbar MN progenitors and no sacral MN progenitors were born earlier (during the E6-E10 period, Fig. 4C). We also found that 24-31% of Olig2^+^ MN progenitors were not labeled by TAM injection at E6, which may have been caused by the contribution of non-NMP-derived NSCs at these caudal levels, or by the insufficient penetration of our genetic manipulations. Much lower contributions of E6- and NMP-derived MN progenitors to brachial and thoracic levels were observed (Fig. S1A, B), which is consistent with the current idea that NMP-derived NSCs are gradually added to the caudal spinal cord during axial elongation. Finally, we addressed NMP-derived NSC differentiation. We found that GFP^+^ cells were observed in dorsal neurons with midline-crossing axonal projections, in ventral MNs, and even in the dorsal root ganglia (DRG) (Fig. 4D). Double labeling with markers for MNs (Isl1/2 or NeuN; Figure 4E, Fig. S1C), or sensory neurons (Brn3a, Fig. 4F, G) confirmed their identities, indicating that NMP progeny contributes to the sensory ganglion neurons in the caudal somatic spinal cord and peripheral nervous system (PNS).

**Figure 4.**
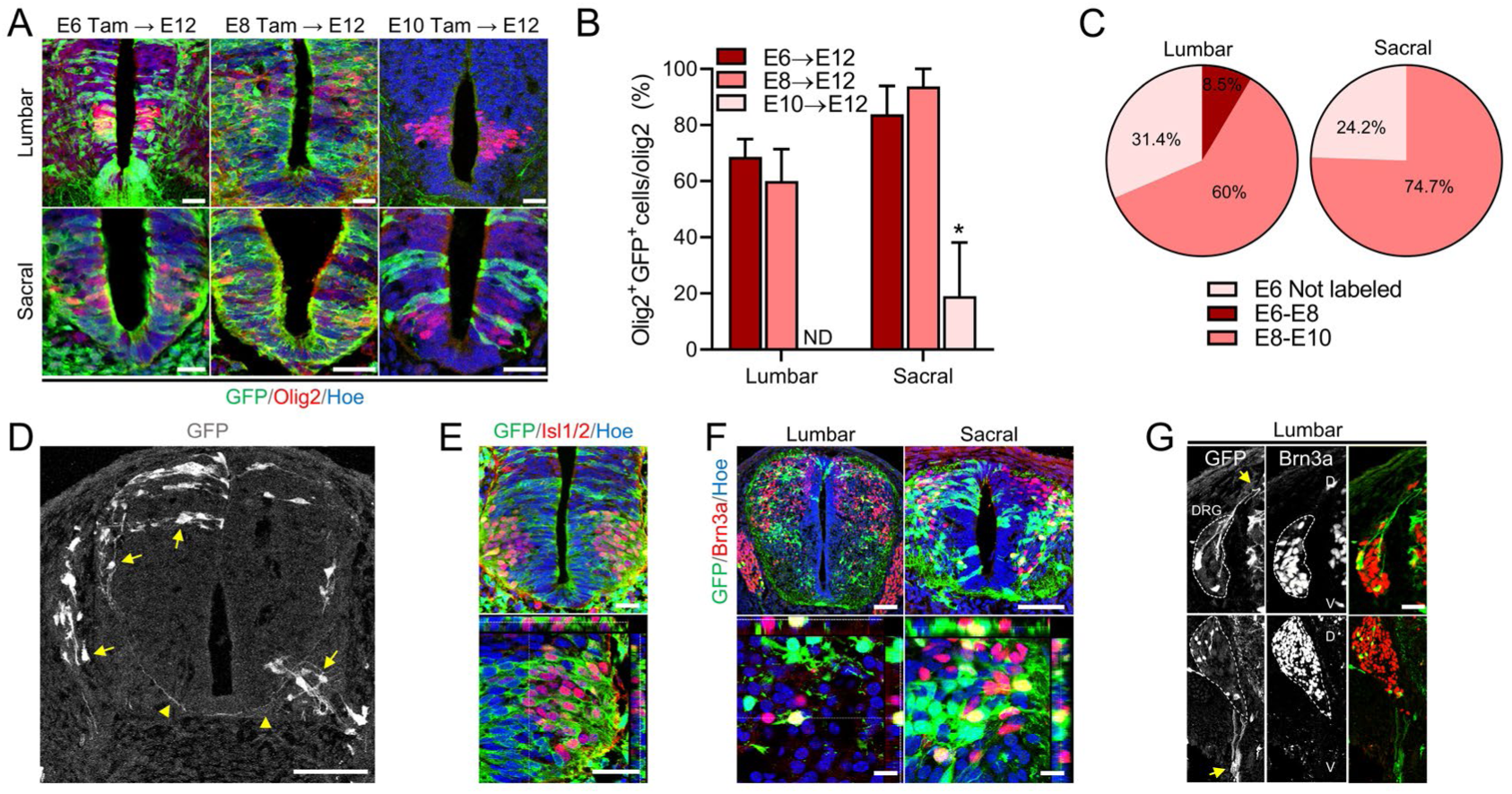
Identification of the progeny of motoneurons and sensory neurons derived from neuromesodermal progenitors. (A) Transverse sections through the lumbar and sacral levels of TCreER:Rosa-EGFP embryos treated with tamoxifen independently at E6, E8, and E10 and sacrificed at E12. Sections are immunostained for Olig2 (Red) and GFP (Green) and counterstained with Hoechst 33342 (Blue). Scale bar = 30 µm. (B) Quantification of GFP^+^ cells expressing Olig2 among the total number of Olig2^+^ cells in lumbar and sacral tissues that are shown in panel (A). Data are presented as mean ± standard deviation. ^*^ P<0.05 One Way ANOVA. (C) Pie chart showing the proportion of GFP^+^ cells expressing Olig2 that are born in E8 and E10 in the lumbar and sacral spinal cord. The proportion is calculated by subtracting the percentages of GFP^+^ and Olig2^+^ cells labeled at E6 from those labeled at E8, and by subtracting the percentages of GFP^+^ and Olig2^+^ cells labeled at E8 from those labeled at E10. All embryos were examined at E12. Proportion of GFP^-^ cells expressing Olig2^+^ are considered not labeled. (D) Lumbar neural tube treated with tamoxifen at E10 and sacrificed at E12; the white color highlights the distribution of GFP^+^ cells. The yellow arrows point to the sensory neurons within the dorsal and DRG domains, while the ventral arrow points to the MNs within the motor column. Arrow heads point to the pre-crossing and post-crossing axons of the commissural neurons. Scale bar = 100 µm. (E) Transverse section through the lumbar level of a TCreER:Rosa-EGFP embryo treated with tamoxifen at E8 and sacrificed at E12 and immunostained for Isl1/2 (Red) and GFP (Green); the bottom panel is an orthogonal optical section. The section was counterstained with Hoechst 33342 (Blue). Scale bar = 30 µm. (F) Lumbar and sacral cross-sections treated with tamoxifen at E10 and harvested at E14. Sections are immunostained for Brn3a (Red) and GFP (Green) and counterstained with Hoechst 33342 (Blue). The bottom panel is a magnified orthogonal optical section. Scale bar = 70 µm, scale bar of orthogonal images = 20 µm. (G) Magnified images of the lumbar DRG of embryos treated with tamoxifen at E10 and sacrificed at E12 (top panel) and E14 (bottom panel). Arrowheads point to the nerve fiber innervation to the dorsal spinal cord at E12 and the ventral innervation of peripheral nerve fibers at E14. Dotted lines mark the morphology of the DRG. D represents dorsal. V represents ventral. All sections were stained for GFP (Green) and Brn3a (Red). Scale bar = 50 µm.

## DISCUSSION

In this study, we discovered that mouse NMPs localize at the caudal neural tube during the neural tube elongation period, where they contribute to posterior neurulation and elongation. On the other hand, chick NMPs were restricted to tail-bud tissue but not to the neural tube. The differences in NMP distribution patterns closely correlate with distinct features of embryonic neurulation processes of the mouse and chick. For instance, a junctional neurulation zone is absent in the mouse due to the continuity of primary neurulation and neural tube elongation, whereas a chick junctional zone is morphologically present. Using an *in vivo* lineage tracing approach, we mapped and illustrated the contribution of NMPs to mouse spinal neural tube formation and found that it gradually increased along the AP axis of the spinal cord, during the axial elongation of the neural tube. We also discovered a tendency for NMP-derived NSCs to contribute to the different regions at the lumbar level of the spinal cord, in a DV gradient, depending on their birthdates. These features are associated with atypical neural tube elongation in caudal neural tube morphogenesis in mice.

Neural folding and neural elongation are different with respect to morphogenetic processes, and the fundamental aspects of this difference appear to be conserved across species. The elongation of the neural tube during development progresses craniocaudally, with the cranial domain developmentally more mature than the caudal domain. Upon fusion of the elongating neural tube with the primary neural tube, a junctional zone known as junctional neurulation is formed and is clearly present in many species, including the chick (Dady et al., 2014). However, other species, such as zebrafish and Xenopus, display no junctional neurulation because the progress of neurulation appears to be continuous with the process of neural tube elongating (Lowery and Sive, 2004). In the mouse, it has also been proposed that a junctional neurulation zone is absent (Schoenwolf, 1984). Our observation that NMPs reside within the caudal neural tube appears to conform to the atypical neural tube elongation and the absence of junctional zone in mice. NMPs are present in the caudal part of the body and contribute to axial elongation; it is believed that they are responsible for caudal spinal cord elongation, through localization at the tail-bud of the chick, mouse, human, and zebrafish (Martin and Kimelman, 2012; Olivera-Martinez et al., 2012; Tsakiridis et al., 2014). Our examination of the spatiotemporal distribution of NMPs at the later developmental phases identified Sox2 and BraT double-positive cells in the mouse caudal neural tube at E10-E11, but with only marginal occurrence in the tail bud. Thus, it appeared that these NMPs within the neural tube differentiated into definite NSCs with the loss of BraT expression. Recently, it was reported that chick caudal NMPs are spatially and functionally segregated, depending on the ability to produce neural vs. mesodermal cells and the placement of neural-lineage restricted cells at the anterior domain (Kawachi et al., 2020). Collectively, although the location of NMPs and detailed caudal morphogenesis are different in the mouse and chick, the fundamental molecular programs for caudal neural tube elongation appear to be conserved between species.

It has previously been reported that cells originated from NMPs are detected within the primary neural tube regions (Gouti et al., 2014; Henrique et al., 2015). Accordingly, *in vitro* differentiation of NMP for 8 days upon activation by retinoic acid led to an increase in the MN progenitor marker Olig2, with posterior identity determined by Hox gene expression. Differentiating cells further adopted neuronal morphology and expressed Tuj1 (Gouti et al., 2014). In addition, 14 days of NMP differentiation under conditioned media favoring the generation of sympathetic neurons generated peripheral neuron-specific intermediate filaments, which express tyrosine hydroxylase, dopamine beta hydroxylase, and peripherin in over 85% of cells (Kirino et al., 2018). In order to study the lineage of NMPs during axial elongation, studies involving dye labeling and cell grafting have been conducted (Kawachi et al., 2020; Martin and Kimelman, 2012). However, these approaches are often imprecise due to the close proximity between various cell types producing various cell mixtures during development of the embryo. To overcome this limitation, we took advantage of TCreERT2 and crossed the strain with a Rosa-EGFP reporter mouse to trace all lineages of BraT^+^ cells emerging from tail- bud tissue (Anderson et al., 2013). We found that an increasing proportion of spinal cord domains in the anteroposterior axis are derived from NMPs, suggesting a co-contribution of NMPs and neuroectodermal progeny in building-up the neural tube. We also determined that NSCs derived from NMPs generate ventral domains first and dorsal domains later in the lumbar neural tube. This is consistent with previous studies which adopted clonal and homotopic graft experiments during early phases of spinal cord development and observed that the majority of early-produced NSCs were restricted to the ventral half of the neural tube (Cambray and Wilson, 2007; Forlani et al., 2003). The DV positions of these clusters are fixed and conserved among animals (Hollyday and Jacobson, 1990). Accordingly, we found that these NMP-derived NSCs properly produced mature neurons populating the spinal cord. We also found that NMP-derived NSCs contributed to sensory neurons in the DRG, which are generated by border cell-derived neural crest cells (Kasemeier-Kulesa et al., 2005). This is consistent with a recent study demonstrating the production of DRG cells from human pluripotent stem cells (hPSCs) via induction of NMPs (Frith et al., 2018), indicating that NMPs exhibit a potential to produce neural lineage cells which contribute to the CNS and PNS.

Despite the significant insights gleaned about the *in vivo* neurobiological patterns of NMPs and their progeny, there remains a surprisingly large gap of knowledge. In-depth *in vivo* studies and cell differentiation experiments in NMP culture *in vitro* based on our current observations will be complementary methods to address how caudal neural tissues are produced and organized in detail.

## MATERIALS AND METHODS

### Animals and embryos

For transgenic mice, the TCreER2 transgenic mouse line used in this work has been described previously (Anderson et al., 2013). TCreER2 transgenic and Rosa-EGFP reporter mice were purchased from Jackson ImmunoResearch Laboratories. TCreER males were crossed with Rosa-EGFP females to generate TcreER2:Rosa-EGFP transgenic males, where the *Rosa26-LacZ* allele is inserted into the *BraT* promoter. In TcreER2:Rosa-EGFP mice, Bra T cells express GFP upon recombination after tamoxifen treatment of pregnant females at a concentration of 0.175 mg/g. Rosa-EGFP females were then back crossed to TcreER2:Rosa-EGFP males, and females were separated on the following day, at embryonic day 1 (E1). Tamoxifen was injected into pregnant females at either E5, E6, E8 or E10 to label BraT cells before and after neural tube closure, respectively. For wild-type mice, pregnant female C57BL/6 mice at E8, E10, E11, and E12 were purchased from DAEHAN BIOLINL. For chick embryos, eggs were purchased from a commercial supplier (Pulmuone, South Korea) and incubated in a turning incubator at 38 °C with relative humidity at 65%. Following incubation, chick embryos at Hamburger-Hamilton (HH) stages 9, 12, and 28 were harvested. The embryos were fixed in 4% paraformaldehyde (PFA) solution overnight. All experiments were carried out in accordance with the ethical guidelines of Korea University and with the approval of the Animal Care and Use Committee of Korea University (KOREA-2016-0125).

### Immunohistochemistry

Tissue processing was performed as described in (Lee et al., 2019) and immunohistochemistry (IHC) was performed as described in (Kim et al., 2017). In brief, tissues were fixed with 4% PFA in 1x phosphate buffer saline (PBS) overnight at 4 °C, followed by washing several times with PBS before sectioning. Fixed tissues were then immersed in 30% sucrose in PBS at 4 °C before being embedded in a solution containing at 3:2 ratio Optimal Cutting Temperature (O.C.T) and 30% sucrose on dry ice. Embedded tissues were then sectioned serially to 14-µM thickness and collected onto New Silane III-coated slides (Muto Pure Chemicals Co. Ltd, 5118-20F). For IHC, sectioned samples were washed three times for 10 minutes at room temperature (RT) before blocking for 1 hour with a solution containing 3% bovine serum albumin and 0.1% triton X-100 in PBS. Primary antibodies, including rabbit anti- Sox2 (1:500; Millipore, AB5603), goat anti-Brachyury T (1:500; R&D systems, AF2085), chick anti-GFP (1:2000; Abcam, ab13970), mouse anti-Isl1/2 (1:200; DSHB, 39.4D5), rabbit anti-Olig2 (1:500; IBL, JP18953), mouse anti-NeuN (1:500; Millipore, MAB377), and goat anti-Brn3a (1:500; Santa Cruz, sc-31984) were added overnight at 4 °C before washing three times with PBS for 10 minutes each at RT. Tissues were then incubated with respective secondary antibodies for 1 hour at RT before mounting and imaging. All tissues and cells were counterstained with Hoechst 33342 from Invitrogen. All images were acquired using confocal microscopy (Leica TCS SP8).

### Whole-mount immunostaining and 3D imaging

Tissue processing was performed as described in (Lee et al., 2020), and whole-mount IHC was performed as described in (Shaker et al., 2015). In brief, dissected tissues were fixed in 2% PFA in PBS for 20 minutes on ice. Fixed embryos were then washed three times for 10 minutes each on ice. Tissues were then dehydrated in 50% methanol in PBS for 10 minutes and then in 100% methanol, two times for 10 minutes on ice. Tissues were either stored at −20 °C or rehydrated in PBST before blocking with 10% BSA in PBST overnight at RT. Primary antibodies (listed above) were diluted in PBST containing 10% BSA and added to blocked tissue for two days at 4 °C. Tissues were then washed three times with PBST for 20 minutes at 4 °C before incubation with secondary antibodies for two days at 4 °C. Tissues were washed again with PBST for 20 minutes at 4 °C before mounting and imaging. All steps were performed using a rocker, with gentle rocking for approximately 15 seconds so as to complete one full rocking motion. All images were captured with a Leica TCS SP8 confocal microscope.

### Imaging, quantification and analysis

Acquired digital images were prepared in Photoshop with adjustment for color, magnification, brightness, and contrast applied equally to images under comparison. For quantitation of cell percentages, total numbers of positive cells were counted manually. Comparisons of lineage-traced cell fates were presented as percent of each sample/tissue section, and multiple samples were used to calculate averages and standard deviations, and for statistical comparisons with one-way ANOVA or unpaired Student’s t-test.

## Statistical analysis

Statistical analyses were performed using one-way ANOVA or unpaired Student’s t-test. All analyses were carried out using the GraphPad Prism 8 software, and the results were presented as mean ± standard deviation. P-values < 0.05 were considered statistically significant.

## Author contributions

Conceptualization: M.R.S., W.S.; Methodology: M.R.S., J.H.L., K.H.K., V.J.K., J.Y.K., J.Y.L., W.S.; Validation: M.R.S., J.Y.L., W.S.; Formal analysis: M.R.S., W.S.; Investigation: M.R.S., J.Y.L., W.S.; Resources: J.Y.L., W.S.; Data curation: M.R.S., J.H.L.; Writing - original draft: M.R.S., W.S.; Writing - review & editing: M.R.S., J.Y.L., W.S.; Visualization: M.R.S.; Supervision: W.S.; Project administration: W.S.; Funding acquisition: J.Y.L., W.S.

## Supplementary Figure

**Figure S1.**
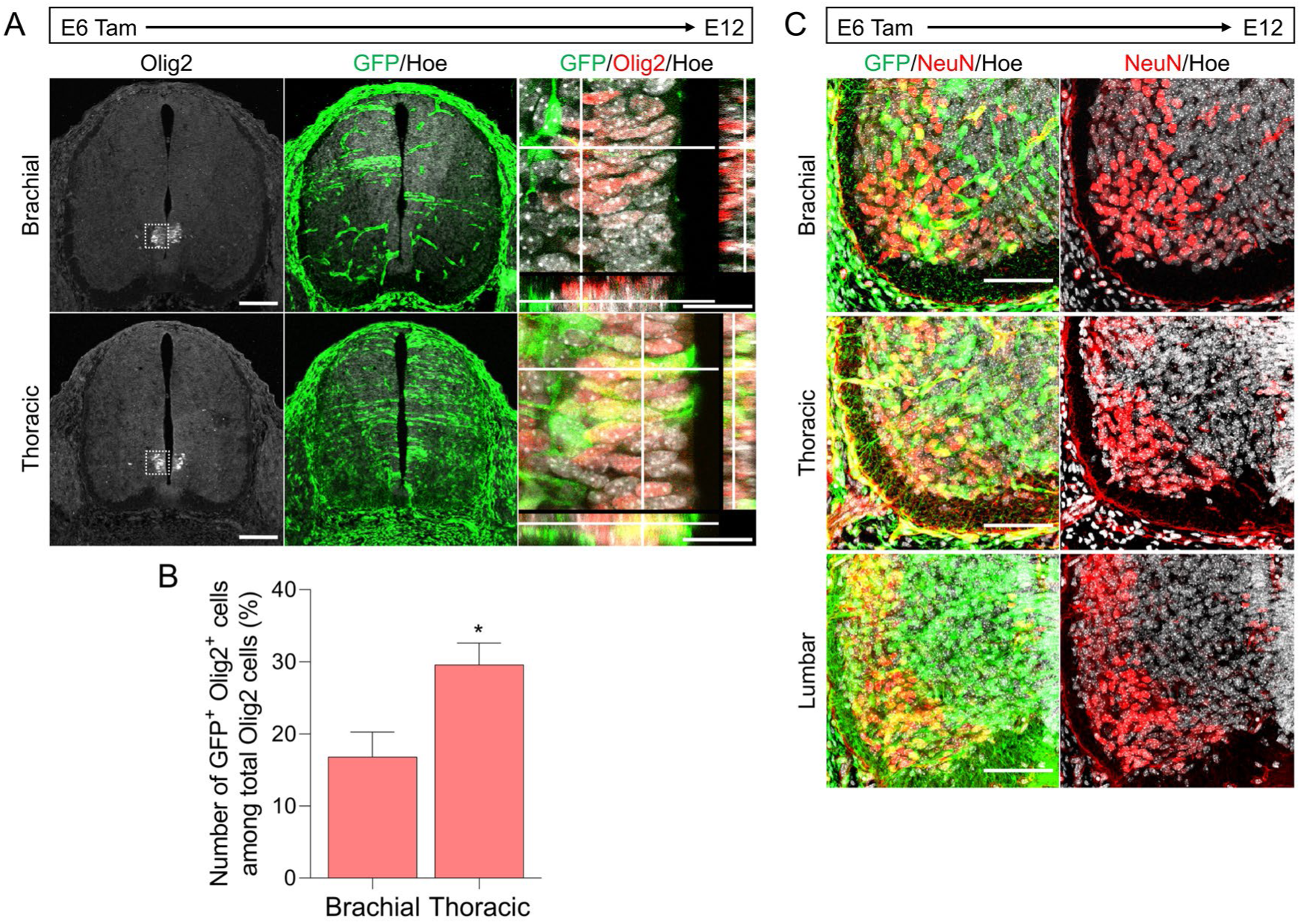
Distribution of the progeny of progenitors and mature motoneurons derived from neuromesodermal progenitors. (A) Transverse sections through the brachial and thoracic levels of TCreER:Rosa-EGFP embryos treated with tamoxifen at E6 and sacrificed at E12. Sections were immunostained for Olig2 (Red) and GFP (Green) and counterstained with Hoechst 33342 (Grey). The right panel is a magnified orthogonal optical section. Scale bar = 30 µm. (B) Quantification of GFP^+^ cells expressing Olig2 among the total number of Olig2^+^ cells observed in the brachial and thoracic tissues that are depicted in panel (A). Data are presented as mean ± standard deviation. ^*^ P<0.01 Paired t-test. (C) Transverse sections through the brachial, thoracic and lumbar levels of the embryonic spinal cord of a TCreER:Rosa-EGFP mouse treated with tamoxifen at E6 and sacrificed at E12. Sectioned tissues were immunostained for NeuN (Red) and GFP (Green) and counterstained with Hoechst 33342 (Grey). Scale bar = 30 µm.

